# Photopatterned spatiotemporal organisation and *in situ* differentiation of 3D human cortical networks

**DOI:** 10.64898/2026.07.08.737264

**Authors:** Siyuan Dong, David Weyland, Hossein Heidari

## Abstract

Modelling human cortical microcircuitry *in vitro* requires platforms that recapitulate both the compositional complexity and spatial architecture of developing neural tissue. Current organoid and assembloid models often rely on the bulk fusion of pre-differentiated, region-specific cells, lacking the capacity for emergent spatial co-differentiation and microenvironment-driven multiscale organisation. There is also a lack of neural and neuronal-glial models with photo-architectured network geometries. To address these limitations, we present a volumetric *in situ* differentiation system using a triculture of precision reprogrammed human iPSC-derived glutamatergic neurons, GABAergic neurons and astrocytes embedded throughout ultra-soft photocrosslinkable hydrogel microenvironments. The deterministic and spatially controlled method allows us to engineer macro-scale, interconnected human neural networks directly onto functional microelectrode array interfaces using projection photopatterning for high-throughput screening. Unlike fusion-based organoids and assembloids, our platform enables simultaneous, spatially distributed lineage differentiation and maturation, and extensive topography-guided neurite outgrowth bridging localised cellular hubs to recapitulate various aspects of neurodevelopmental patterning and synaptic integration in 3D. The model enables topographic patterning of neuronal-glial networks as well as 3D cell-embedded bioprinting with the developed triculture system. Both modes of cellular growth are studied and demonstrated here. Longitudinal electrophysiological tracking over a month of culture reveals a transition from immature, quiescent states to asynchronous, information-dense microcircuits characterised by an expanded state-space manifold and physiological excitatory-inhibitory balance. By replicating the mechanics of native brain parenchyma, the model presents a highly reproducible, scalable and flexible platform for the study of cortical microcircuitry development, neurodegenerative decline, and inter-regional network assembly.

## 1. Introduction

The human nervous system is an intricate and highly organised network of neurons and supporting cells that enable communication and information transfer across the entire body. Functionally, it is divided into the central nervous system (CNS) and the peripheral nervous system (PNS), which together regulate essential physiological processes and sustain normal life activities^1^. Perturbations of neuronal connectivity or progressive neuronal loss can disrupt signal transmission, leading to neurodegenerative disorders which are characterised by chronic and progressive deterioration of network structure and function^2,3^. The accelerating prevalence of such disorders including dementia and Parkinsons demands elevated clinically-relevant research priority^4,5^. There is a need for improved *in vitro* models that can more accurately and controllably capture their development and progression mechanisms to help accelerate therapeutic discovery and development.

Traditionally, these mechanisms have been studied using animal models followed by two-dimensional tissue culture systems. This approach has significant limitations with animal models often failing to reproduce the full spectrum of human disease, particularly with respect to genetic heterogeneity, aging-related processes, and species-specific differences in neuronal circuitry. As a result, many preclinical findings have not translated into clinical therapies^6–8^. In addition, two-dimensional models cannot fully mimic the complex 3D environment, topographies, extracellular matrix (ECM) interactions, and spatial cellular diversity required for a physiologically relevant human brain model^9–11^. To address these translational gaps and limitations, 3D *in vitro* models have been sought which would allow us to engineer physiologically relevant neural microenvironments. Natural and synthetic hydrogels with tuneable mechanical properties and porosity^12–14^, have been developed which could support key cellular behaviours including neurite outgrowth, cell mobility and motility, and localised connectivity, which determine the maturation and functional integration of engineered neural tissues^15,16^. However, there is very limited work demonstrating successful 3D differentiation and growth of co-culture iPSC-derived human neuronal-glial tissues. There is also a lack of neural models fabricated with techniques such as extrusion, droplet deposition, layer-by-layer and volumetric bioprinting to enable spatially controlled cellular organisation^17,18^.

While microfluidic and droplet-based neuroglial and neurovascular assembloid models successfully incorporate neurons, astrocytes and endothelial cells to demonstrate functional features (synaptic activity, lumen formation, and partial barrier behaviour)^19–21^, the core idea of multi-region and multi-lineage assembloids which relies on fusing high-density region-specified organoids after lineage commitment, limits their ability to mimic *de novo* emergence of network composition and balancing of excitatory–inhibitory and astrocytic modulation typical of early human corticogenesis. As a result, structural organisation is frequently achieved without fully reproducing the dynamic and functional cellular coordination observed in native neurovascular development.

An additional limitation of current neuronal-glial organoids and hydrogel-based culture systems is the lack of spatially controlled network tissue structure, architecture and composition. Although three-dimensional ECMs support neuronal growth and self-organisation at pellet densities, cellular assemblies often emerge randomly with limited control over tissue geometry, connectivity patterns or network architecture. On the contrary, the developing human brain presents highly organised structure patterning across multiple spatial scales, which contribute to cellular localisation, neurite guidance and circuit formation. There is a clear need for engineered topographies and microenvironment. However, the application of biofabrication techniques to human neuronal-glial systems remains unexplored, especially in a platform that would combine spatial patterning and *in situ* differentiation with electrophysiological tracking interfaces.

Here, we describe a photopatterned *in situ* differentiation and neuronal tissue growth platform for neuroglial co-organisation. To demonstrate this concept, we introduce a novel scalable triculture of human iPSC–derived glutamatergic, GABAergic, and astroglial progenitors patterned with engineered photoactive hydrogel matrices. We demonstrate physiological relevance and reliability of the model and its capability to mimic critical aspects of neural tissue development, and exhibit specific functional indicators previously limited to 2D models^22–24^.

## 2. Results and discussion

We demonstrate *in situ* differentiation and topographical and microenvironmental guidance of neuronal and glial 3D architectures. We show the formation of networks of neuronal-glial organoids from isolated iPSC cells across a map of macro-topographies photo-patterned on top of an array of functional microelectrodes. The proposed approach provides a unique window to spatiotemporal screening, instruction and perturbation of large multi-organoid networks in a way never achievable prior to this with aggregation-based approaches such as hanging droplets and gravitational microwells. The construction of such functional and scalable human neural circuits *in vitro* requires an optimal integration of cell types and densities, extracellular matrix biophysics, and multiscale architecture. The system replicates the traction, compliance, and extracellular cues of native brain parenchyma.

A physiologically relevant, human-derived neuronal-glial triculture system was developed within a biomimetic hydrogel framework. This platform utilises a photocrosslinkable gelatin methacrylate (GelMA) hydrogel at a low concentration of 6% (w/v) crosslinked with lithium phenyl-2,4,6-trimethylbenzoylphosphinate, and seeded with a well-defined and highly reproducible triculture of human induced pluripotent stem cell (hiPSC)-derived cells obtained through precision cellular reprogramming. The model incorporates excitatory Glutamatergic neurons, inhibitory GABAergic neurons, and supporting astrocytes, establishing a representation of the central nervous system (CNS) microenvironment. This platform is presented with two distinct engineered configurations. The first configuration is a topographic model where photopatterned hydrogel topographies serve as structured 3D surfaces for topography-guided network formation from surface-seeded neuronal-glial populations^25^. The second configuration is a 3D cell-embedded model where neuronal-glial populations are directly embedded in photopatterned hydrogels to facilitate volumetric self-organisation, migration, and neurite outgrowth (Fig. 1G).

**Figure 1.**
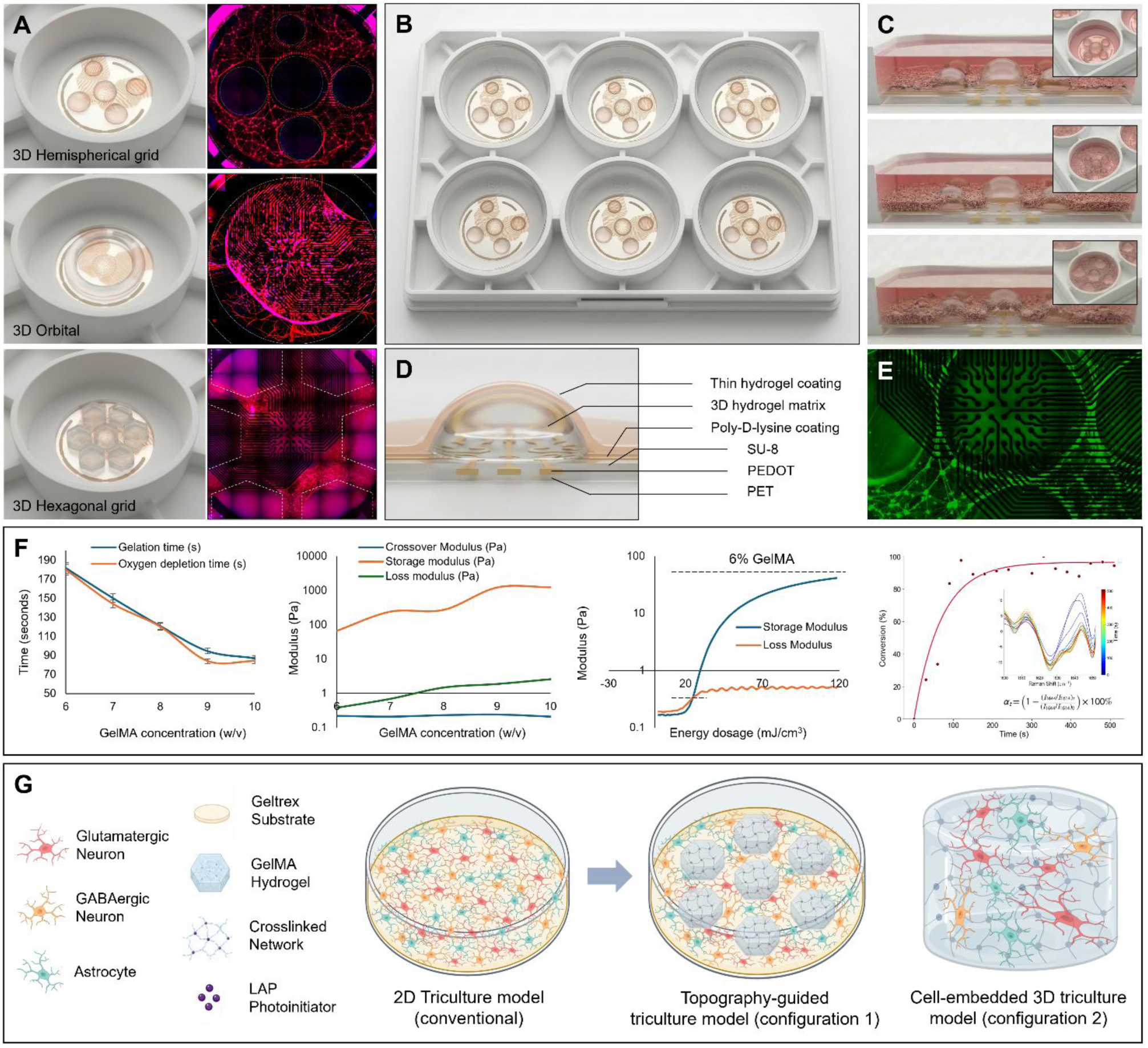
Engineered photopatterned hydrogel platforms integrated with MEAs for cortical network modelling: (A) topographies fabricated on MEA substrates in multiwell plates (B), together with corresponding live cell tubulin red staining demonstrating successful neuronal-glial network formation within each geometry; (C) illustration of post-print topography-guided neuronal-glial development; (D) device structure for multiwell tissue models; (E) fully-developed neuronal-glial network morphology; (F) physicochemical characterisation of photoactive GelMA hydrogels; and (G) illustration of demonstrated tissue configurations.

### 2.1. Photopatterned microenvironments with integrated microelectrode arrays for cortical modelling

One of the major challenges in neural tissue engineering is the ability to establish spatially controllable neural tissues. The formation of brain organoids and assembloids in current models depends on bulk and random self-organisation processes, resulting in significant variability in architecture, network morphology and functional maturation. To overcome these limitations, we developed a photopatterned hydrogel platform integrated into microelectrode arrays as a reproducible engineered neuronal-glial platform with defined geometries and real-time functional measurement. Various macroscale topographical configurations were printed using light-based projection biofabrication, including 3D hemispherical and hexagonal grids, and orbital structures (Figs. 1A and B). Configurations were designed to provide different levels of confinement and interconnection, enabling us to investigate the influence on neuronal-glial self-organisation *in vitro*. The hemispherical grid forms individual three-dimensional niches that constrain initial cell distribution and network formation. On the contrary, the orbital and hexagonal structures present more continuous topographical features which can guide cellular migration and neurite outgrowth over extended distances (Figs. 1C, E and G), allowing us to investigate how complex configurations affect emergent neuronal-glial organisation. Large-scale organisation of stable neuronal-glial networks and extensive neurite connections observed throughout the patterned regions, demonstrates that the engineered microenvironments support neuronal survival, process extension, and network formation. The final spatial distribution of neuronal processes exhibits a close correspondence with the underlying hydrogel geometries, suggesting that macro-scale design contributes to the organisation of neuronal-glial assemblies.

The human brain microenvironment provides an extremely soft and degradable matrix, with ECM stiffness on the order of only a few hundred Pascals in development and up to a few kilopascals in adulthood. Neurons are highly mechanosensitive thriving in matrices that match this compliance. In fact, sub-kPa substrates optimise neuronal differentiation, while significantly stiffer or softer environments can reduce the yield of neuronal network formation^26^. We target the early neuronal-glial development and are therefore interested in the few hundred Pascal domain. Other 3D culture models have deliberately tuned hydrogel stiffness to ∼1 kPa to mimic the brain^27^. Fully converted GelMA hydrogels at 6-7% (w/v) concentration, yield storage moduli of 100-300 Pa (Figs. 1D and F), well within the developing brain’s native mechanical range^28^. Such ultra-soft GelMA gels provide the compliant microenvironment that neurons require for extending neurites and forming networks. Matrices such as methacrylated hyaluronic acid (HAMA) were also implemented as predominant components of brain ECM^29,30^. HAMA was removed from our system due to the major stiffness contribution even at a concentration of 0.25% (w/v) which was the lowest stable concentration over multiple weeks (Addition of a 0.25% (w/v) of HAMA pushed the original storage moduli of 6% GelMA from 100 Pa to well over 1000 Pa). Hydrogel matrices based on GelMA have been widely used for 3D cultures due to their tuneability^31^. The GelMA 6% was ultimately chosen as a matrix that could maintain construct integrity over multiple weeks, while encouraging neurite outgrowth and supporting the triculture expansion in 3D without provoking excessive astroglial reactivity that can emerge in overly stiff gels. The synthesised GelMA 6% with a >95% degree of substitution was able to support the triculture of GABAergic, Glutamatergic neurons, and Astrocytes in the following studies. The *in situ* monitoring of the acrylate photopolymerisation kinetics yielded an initial rapid alteration in the normalised peak ratio within the first 90 s of exposure, reaching a calculated conversion variation of 23.63% (Fig. 1F). Beyond this initial regime, the system transitioned into a steady-state characteristic of a non-reactive spectator environment. At the lowest tested concentration of 6% w/v, the oxygen depletion time and gelation time are at their highest, measuring approximately 180 seconds. As the GelMA concentration increases to 10% (w/v), both parameters decrease significantly, levelling off at approximately 85–90 seconds (Fig. 1F). The close tracking between the oxygen depletion curve and the gelation curve highlights the profound impact of oxygen inhibition on radical photopolymerisation.

The presence of both glutamatergic (excitatory) and GABAergic (inhibitory) neurons is essential for the establishment of functional neural circuits. Glutamatergic neurons provide the primary excitatory drive, while GABAergic interneurons regulate the timing and synchronisation of network activity^32^. In the fully developed networks, the long neurites represent network buses, potentially forming the basis for long-range white matter-like tracts that connect the distant soma clusters^33^. In a healthy human brain, astrocytes play key roles in synaptic pruning, metabolic support, and neurotransmitter homeostasis. However, when cultured on rigid glass or plastic substrates (with mechanical stiffnesses in the gigapascal range), astrocytes undergo reactive astrogliosis. This reactive state is characterised by cellular hypertrophy, pathological upregulation of glial fibrillary acidic protein (GFAP), and the secretion of chondroitin sulphate proteoglycans that actively inhibit neurite extension and damage neuronal health^34^. The developed 6% GelMA hydrogel possesses an elastic modulus tailored to replicate the development mechanics of native brain tissue. This compliant mechanical environment, combined with the biomimetic polypeptide backbone of GelMA, suppresses reactive astrogliosis. Astrocytes atop or within this matrix maintain a physiological, non-reactive phenotype characterised by highly spread morphologies, actively traversing the nets and supporting synaptogenesis and neuronal survival.

### 2.2. Large-area self-organised *in situ* co-differentiation of neuronal-glial networks

To establish a baseline neuronal-glial triculture prior to the introduction of engineered hydrogel topographies, the system was first evaluated on microelectrode arrays under standard two-dimensional culture conditions. Live-cell tubulin staining was used to visualise the development of viable neuronal structures across the recording surface. The whole-well imaging across centimetre-large networks revealed extensive tubulin-labelled cellular structures distributed throughout the massive culture area. The neuronal-glial triculture expands across the entire recording region, generating a continuous network-like architecture. This widespread distribution indicates robust cellular attachment, survival, and growth across Geltrex-coated microelectrode array substrates. The dense meshwork of tubulin-labelled cellular processes throughout, and numerous neurite bundles bridging neighbouring cellular cluster nodes, generate a complex self-emerged network architecture across the substrate.

The widespread presence of these neurite-like processes suggests that neuronal growth was not restricted to isolated regions but instead occurred across the entire culture surface (Fig. 2). Notably, the cellular network distributed evenly across the coated surface in the well, even on the regions containing dense microelectrode routing tracks and contact pads. This demonstrates the substrate compatibility with the triculture system allowing us to record signals for long term monitoring of developing neuroglial networks.

**Figure 2.**
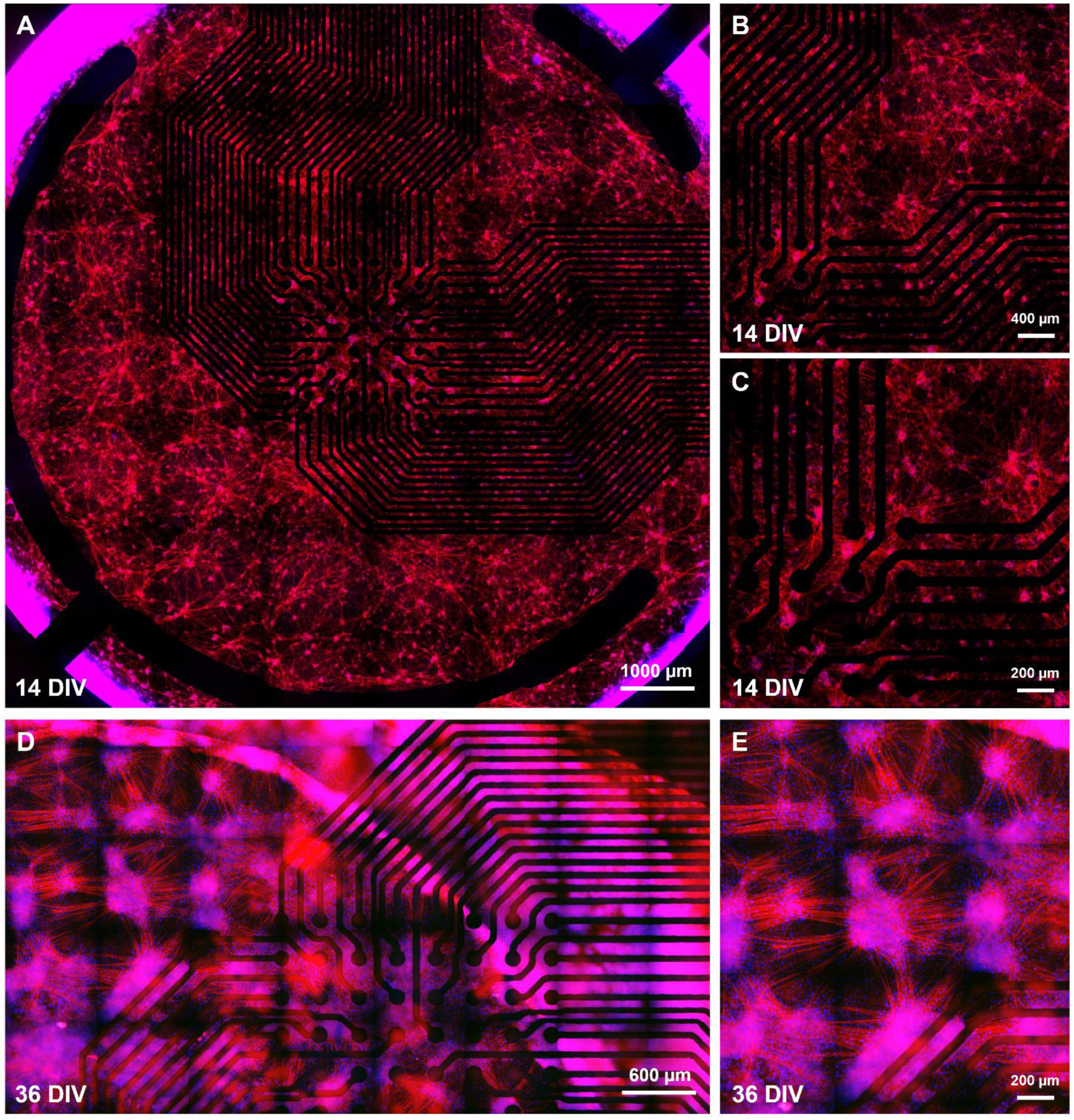
Representative live-cell Tubulin Red staining of the 2D neuronal–glial triculture established on top of a transparent MEA: (A) whole-well imaging demonstrates extensive neurite networks spanning the entire recording area at 14 DIV; (B) and (C) higher-magnification images show dense interconnected neuronal processes distributed across different regions of the culture; (D) fully-matured multi-organoid network spanning a remarkable 10mm wide area with large hubs of over 1000 cells each at 36 DIV; higher magnification images show the massive hubs and large density of thick neurite bundles connecting the hubs.

The distributed network morphology observed in this sparse system differs from aggregation-based neural culture systems, where cells are typically packed into a dense and confined spheroid structure. The widespread distribution of the network of cellular clusters across the microelectrode array substrate, allows us to measure across large recording areas and networks of connected organoids for longitudinal monitoring of development (Fig. 2). The triculture system supports the formation of viable, large area neuronal-glial cortical assemblies on MEA substrates with cellular hubs of thousands of cells in density, and a stable model for tandem investigation of cellular organisation, spatial pattering and network development.

### 2.3. Longitudinal multi-scale functional activity of neuronal-glial networks

The electrophysiological activity of the glial linked assembloid networks (GLANs) was longitudinally and continuously monitored using microelectrode arrays. Morphological integration of the assembloids over the recording electrodes (Fig. 3) was accompanied by a robust, time-dependent escalation in spontaneous network activity (Figs. 3A–C). Analysis of individual GLAN trajectories reveals a highly non-linear increase in the mean firing rate over 36 days of *in vitro* development. This intensification of activity is coupled with an expansion in spatial coverage as indicated by the progressively larger marker sizes (Fig. 3D) demonstrating that as the networks mature, they not only fire at higher frequencies but also recruit a significantly larger proportion of the array into active signalling. Ridgeline plots detailing the probability density of electrode firing rates highlight a critical shift in population dynamics (Fig. 3E). At early time points (e.g., DIV 8–12), the networks exhibit a narrow, low-frequency distribution indicative of a quiescent, immature state. By later stages (e.g., DIV 30–36), the distributions broaden significantly and shift rightward, reflecting the emergence of distinct functional heterogeneity and a wider dynamic range of neuronal activity characteristic of mature microcircuits.

**Figure 3.**
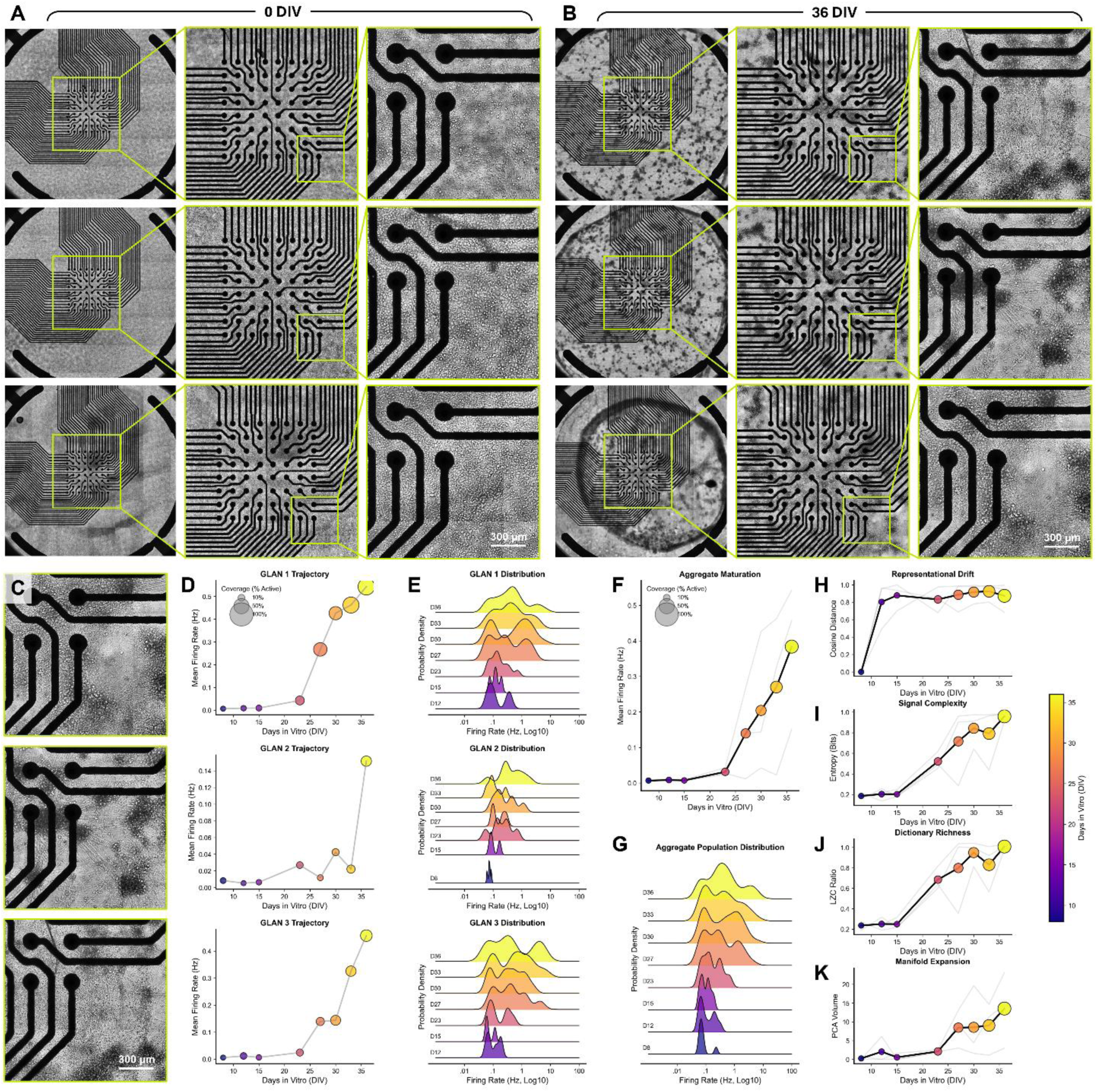
Functional maturation and complex ensemble dynamics of glial-linked assembloid nets (GLANs): (A–C) escalation of spontaneous network activity and morphological integration over time; (D) individual GLAN trajectories showing a non-linear increase in mean firing rate and spatial array recruitment over 36 DIV; (E) firing rate probability densities, shifting from narrow, low-frequency states to broad functional heterogeneity; (F–G) aggregate maturation trajectories demonstrating cross-replicate consistency and population distributions reflecting *in vivo*-like variance; (H) representational drift illustrating rapid initial plasticity followed by network stabilisation around DIV 15; (I–J) escalation of compute capacity, measured by permutation entropy and Lempel-Ziv complexity; and (K) manifold expansion indicating increased dynamic range in mature microcircuits.

To evaluate the reliability of the network developmental trajectory across biological replicates, aggregate maturation metrics were computed. The pooled aggregate trajectory confirms a highly reproducible acceleration in mean firing rate that typically initiates around DIV 20 and culminates in a highly active state by DIV 35, with shaded variance bands indicating tight cross-replicate consistency (Fig. 3F). The corresponding aggregate population distribution corroborates this functional maturation at the macro-level (Fig. 3G). The transition from a unimodal, low-activity peak to a broad, heavy-tailed distribution underscores that the model consistently develops a complex architecture comprising both highly active hub-like regions and sparsely firing zones, mimicking the physiological variance observed in intact *in vivo* neural networks. Beyond first-order firing statistics, the computational capacity and ensemble dynamics of the GLANs were evaluated using advanced topological and information-theoretic metrics. This approach determines whether the *in vitro* activity resembles functionally mature neural processing rather than stochastic or pathologically hypersynchronous firing. Representational drift, quantified via the cosine distance of the population vector relative to its nascent state, provides insight into the temporal stability of the network’s functional identity. The data reveals a biphasic trajectory characterised by a rapid initial divergence that stabilises around DIV 15 (Fig. 3H). This suggests that the network undergoes a period of intense early synaptic plasticity and structural reorganisation akin to developmental critical periods *in vivo* before settling into a highly stable, functional architecture. The stabilisation of the population vector indicates the formation of a robust attractor state, which is a physiological prerequisite for reliable information storage and consistent microcircuit performance.

Concurrently, the computational richness of the network expands dramatically as the assembloid network structurally integrates. Signal complexity, measured through permutation entropy, and dictionary richness, assessed via Lempel-Ziv complexity (LZC), both exhibit robust, monotonic increases over time (Figs. 3I and J). Permutation entropy captures the nonlinear unpredictability of temporal spike trains, while LZC quantifies spatiotemporal signal diversity across the array. The concurrent rise in both metrics demonstrates that the maturing GLAN transitions from emitting simple, redundant, and highly correlated spike patterns which are often associated with immature or seizure-like states, to generating highly variable, information-dense spatiotemporal sequences. This shift highlights the emergence of a sophisticated balance between functional integration and segregation, allowing the microcircuit to maximise its information-carrying capacity.

The network’s overall dynamic range was mapped using manifold expansion, defined by the volume of the state-space traversed by the network following principal component analysis. This geometric metric increases prominently at later developmental stages. An expanded neural manifold directly correlates with an increase in the network’s degrees of freedom; mature GLANs exhibit vastly enlarged dynamic range, allowing them to flexibly explore and sustain a significantly wider array of complex functional microstates compared to early-stage cultures (Fig. 3K). Biologically, this expanded state-space volume mitigates catastrophic interference during stimulus processing and is essential for multidimensional neural computations, confirming that GLANs develop the high-dimensional functional architecture characteristic of intact cortical networks.

### 2.4. Functional biomarker expression in neuronal-glial networks

To validate the neuronal phenotype established within the triculture system, immunofluorescence staining was performed using microtubule-associated protein 2 (MAP2), a well-established marker of mature neurons and dendritic structures, together with nuclear staining. This analysis is conducted to validate neuronal maturation and investigate the spatial distribution of cells throughout the culture, and to complement the live-cell tubulin staining. Confocal imaging showed extensive MAP2 expression across the analysed regions of the neuronal-glial triculture. MAP2-positive neuronal cell bodies and associated dendritic processes were observed throughout the culture, indicating the successful establishment of mature neuronal populations within the triculture system.

MAP2-labelled processes extend between neighbouring cellular regions, in the formation of complex dendritic architectures present in the developing neuronal networks. Confocal images (Fig. 4) demonstrate consistent MAP2 immunoreactivity across the volume, indicating that neuronal morphology was not restricted to a single plane and supporting the widespread distribution of mature neuronal structures across the culture. In addition, MAP2-positive neuronal processes were observed directly in the microelectrode regions, indicating the good compatibility between the developing neuronal network and the MEA substrate critical for electrophysiological signal reliability. Together with the live-cell tubulin imaging presented in section 2.2, these findings demonstrate the successful formation of a biologically relevant neuronal-glial network for subsequent investigations.

**Figure 4.**
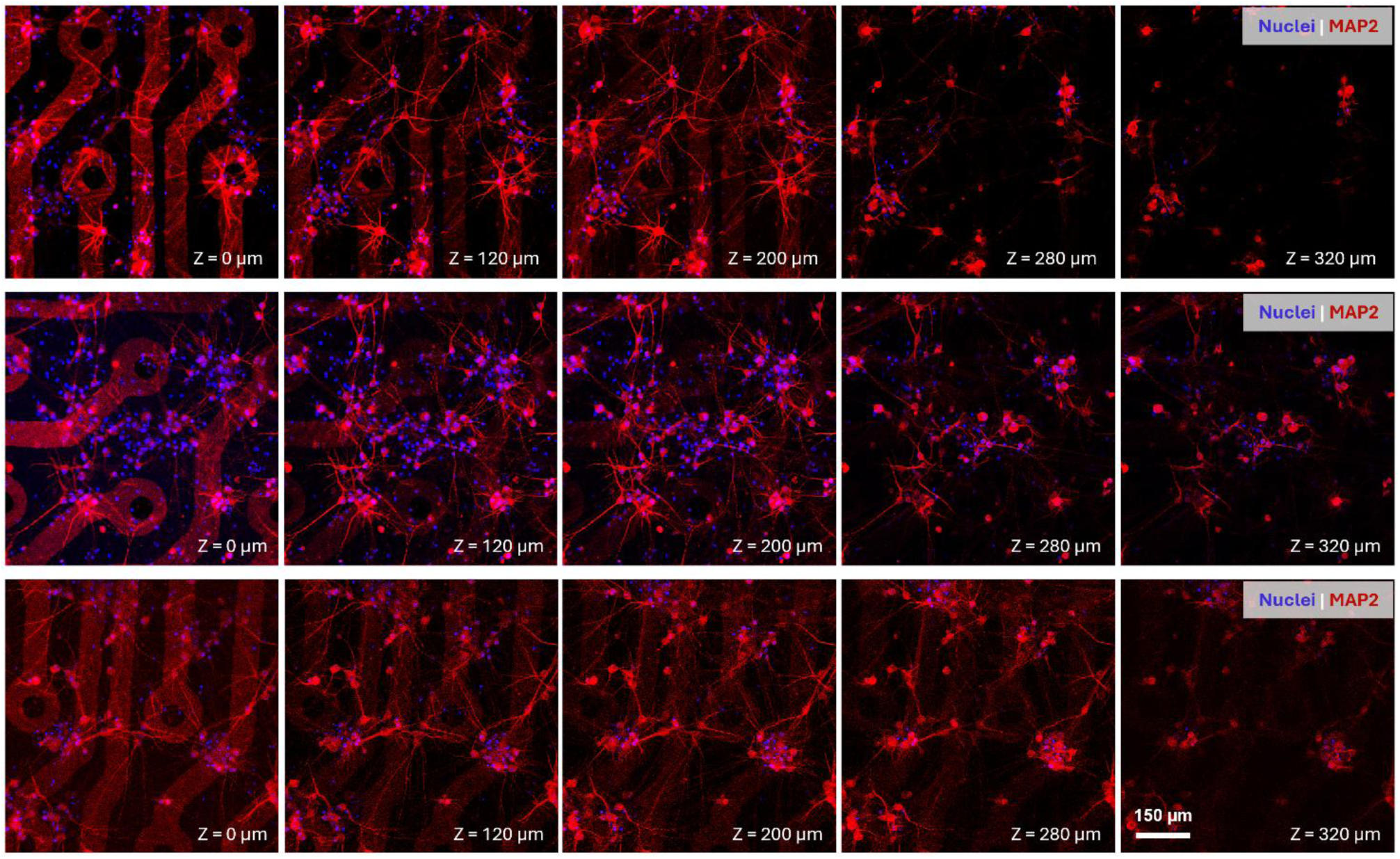
Functional biomarker expression of neuronal–glial networks on transparent microelectrode arrays. Representative confocal z-stack images of MAP2-positive neurons (red) and nuclei (blue) acquired from three representative regions of the 2D neuronal–glial triculture. Images were collected at five imaging depths (0–320 μm), to demonstrate widespread neuronal distribution and persistent MAP2-positive neuronal cell bodies and dendritic processes throughout the culture.

### 2.4. Photopatterned topography-induced neuronal-glial network morphology

With a stable neuronal-glial triculture system established, we investigated whether photopatterned hydrogel topographies could influence cellular organisation and network formation. Live brightfield and phase-contrast imaging was performed over 2-5 weeks with time-lapse recordings every two hours to study progressive cellular reorganisation around the hydrogel structures over hundreds of frames (Fig. 5).

**Figure 5.**
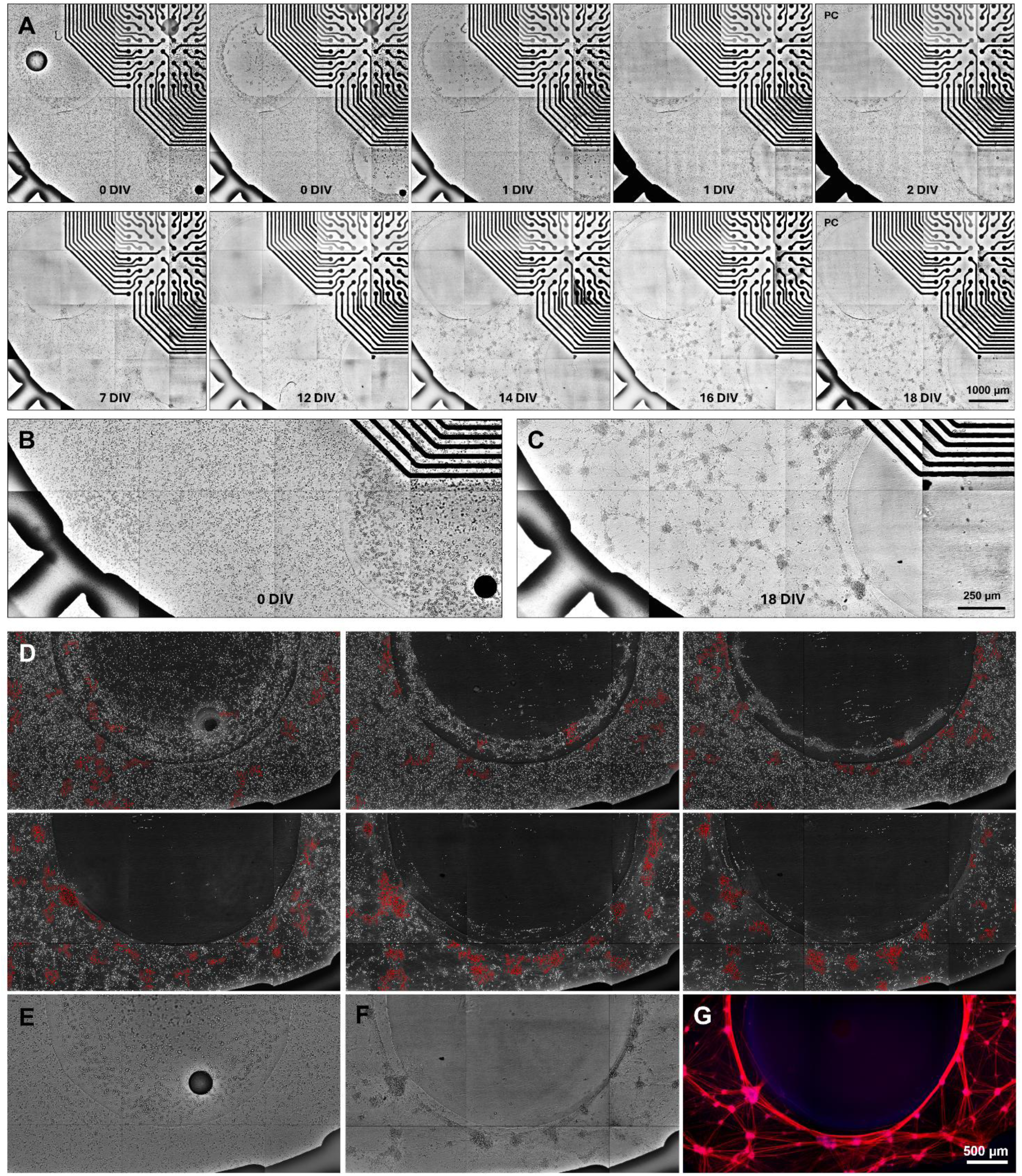
Photopatterned hydrogel topographies guide long-term neuronal–glial self-organisation: (A) representative brightfield images showing the temporal reorganisation of the neuronal–glial triculture proximal and distal to the hemispherical GelMA hydrogel topographies from DIV 0 to DIV 18; (B) and (C) initial and final states across larger area; (D) cells progressively migrated from an initially homogeneous distribution towards the hydrogel interface, where localised cellular clusters gradually form during culture; (E) and (F) ; (G) Tubulin-positive neurites formed continuous interconnected networks while maintaining close spatial association with the engineered topography.

Live imaging revealed a transition from an initially dispersed cellular population to an organised architecture characterised by cellular clustering and extensive neurite outgrowth. By DIV 12, a high density of large clusters (>100 cells) had formed across the culture, interconnected by long neurite projections (Fig. 5D). In the meantime, elongated neurite-like processes extend towards hydrogel regions (Fig. 5G). The hydrogel architecture promotes boundary-associated cluster formation resulting in the emergence of spatially organised neuronal-glial assemblies that were not observed in conventional 2D cultures (Figs. 5C, and E–G).

The hemispherical topography promotes cellular clustering and neurite extension in a unique way, leading to the emergence of spatially organised neuroglial architectures. A patterned distribution is repeatedly observed emerging in close proximity of curved topographies (Figs. 6B and C). This seems to be a result of cells rapidly migrating over the curvature during the first two days. During this topography-facilitated migration, there is an increased likelihood of cells crossing paths and connecting, hence accelerating the cellular organisation and clustering process (Fig. 6A). The 3D hemispherical topographies not only confine but also encourage clustering. Instead of remaining randomly distributed across the substrate, neuronal-glial populations progressively reorganise into regions spatially defined by the hydrogel topographical map.

**Figure 6.**
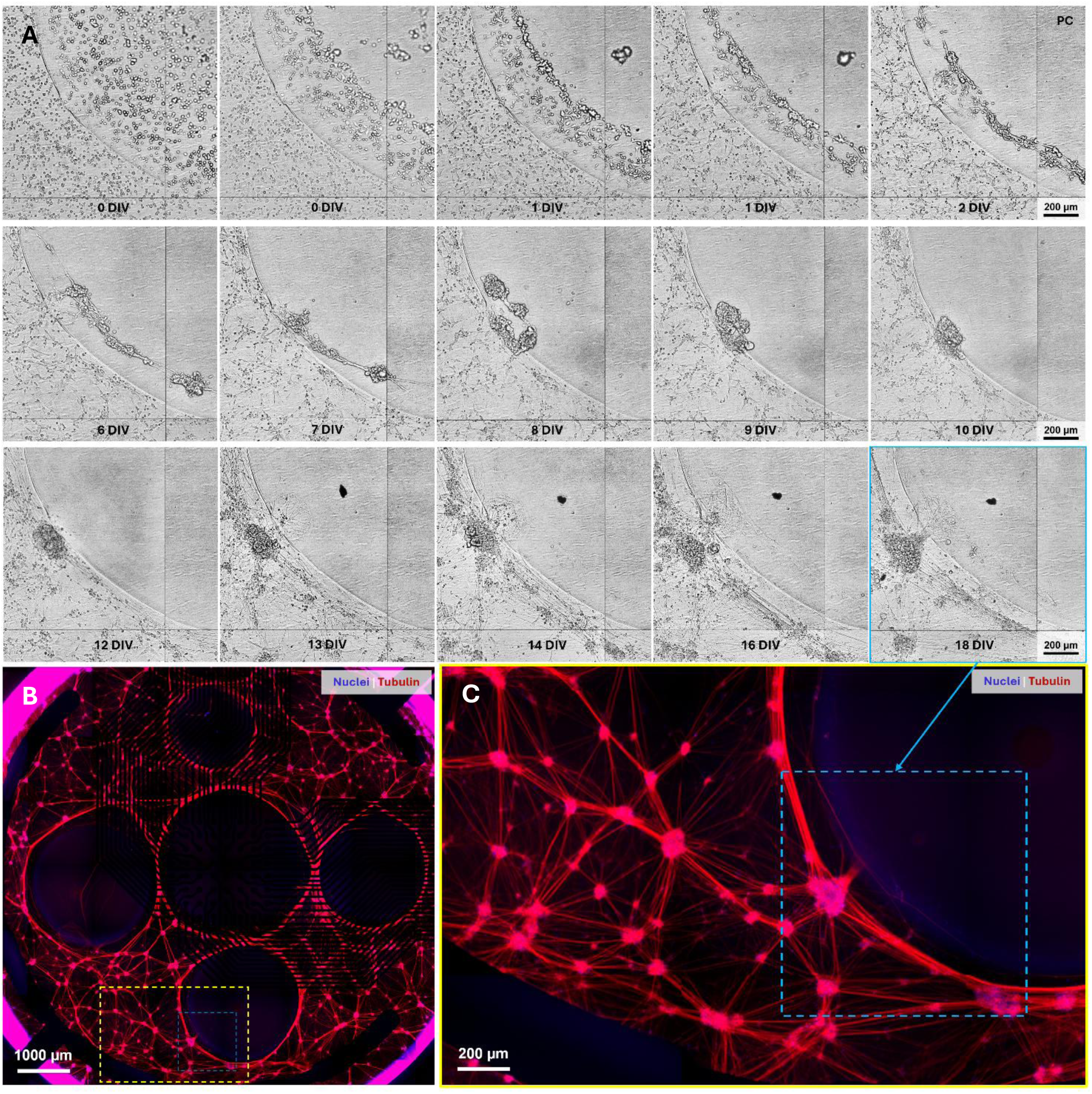
Boundary-associated cell aggregation and the appearance of elongated neurite-like processes extending towards the hydrogel region are observed at later culture stages: (A) brightfield images comparing the cellular distribution at DIV 0 and DIV 18; (B) whole-well live-cell tubulin staining demonstrating large-area neuronal-glial network organisation surrounding hemispherical hydrogel structures; (C) representative higher-magnification images showing dense neurite bundles connecting neighbouring cellular clusters surrounding the hydrogel structures. Similar organisational behaviour is observed across geometries of different diameters, demonstrating reproducible topography-guided neuronal–glial network formation.

Similar observations have been reported in neural tissue engineering studies, where micro-topographical features such as pillars and protrusions influence single cell adhesion, migration, and neurite organisation. However, most of the previous studies have focused on single or few cells in monoculture or biculture systems. In contrast, this platform combines human glutamatergic and GABAergic neurons with astrocytes across a macroscopic photopatterned hydrogel architecture directly integrated with electrode arrays. Rather than simply directing individual cellular behaviours, the topographies influence the collective spatial organisation of developing neuronal-glial populations, providing a unique platform to investigate how engineered tissue architecture contributes to network assembly and maturation.

Live-cell tubulin staining demonstrates that neuronal structures closely followed the imposed hemispherical architecture. Extensive neuronal processes were observed surrounding the hydrogel, forming interconnected network structures outlining the hemispheres. At larger scale, neuronal networks form a continuous architecture across multiple hemispherical areas and maintain a clear spatial association with the patterned topography. Across hemispherical structures of varying diameters from 1.6 to 2.8 mm, similar organisational behaviours were observed indicating that topography-driven cellular patterning is robust across multiple length scales (Figs. 6B and C). Compared with conventional neural cultures, in which cells are typically distributed randomly across the substrate and are difficult to spatially control, the photopatterned millimetre-scale hydrogel topographies promoted reproducible cellular organisation and the formation of defined neuronal-glial target zones.

### 2.5. Neuronal-glial network neurite outgrowth in photopatterned hydrogel microenvironments

Building on the topography-promoted neuronal migration and aggregation at hydrogel boundaries, we next investigate whether alternative photopatterned geometries could further influence neurite outgrowth and neuronal-glial network organisation. To evaluate this, a hexagonal grid architecture with vertical walls was designed to replace the curved edge topography of the hemispherical features and provide interconnected hydrogel channels to enable neurite extension into the engineered microenvironment. Live imaging of the hexagonal grid architecture showed that the patterned hydrogel geometry influences the spatial distribution of the neuronal-glial culture over time. Instead of forming a uniform cell layer, cells progressively organised into patterned regions that corresponded to the geometry of the hydrogel channels. Cellular clusters were observed within the channel-associated regions, indicating that the interconnected grid architecture guided the spatial organisation of the developing culture (Figs. 7A and B). A progressive transition from an initially homogeneous cell distribution at 0 DIV to the gradual formation of localised high-density clusters at the hydrogel boundaries as soon as 14 DIV, was replicated in the hexagonal case.

**Figure 7.**
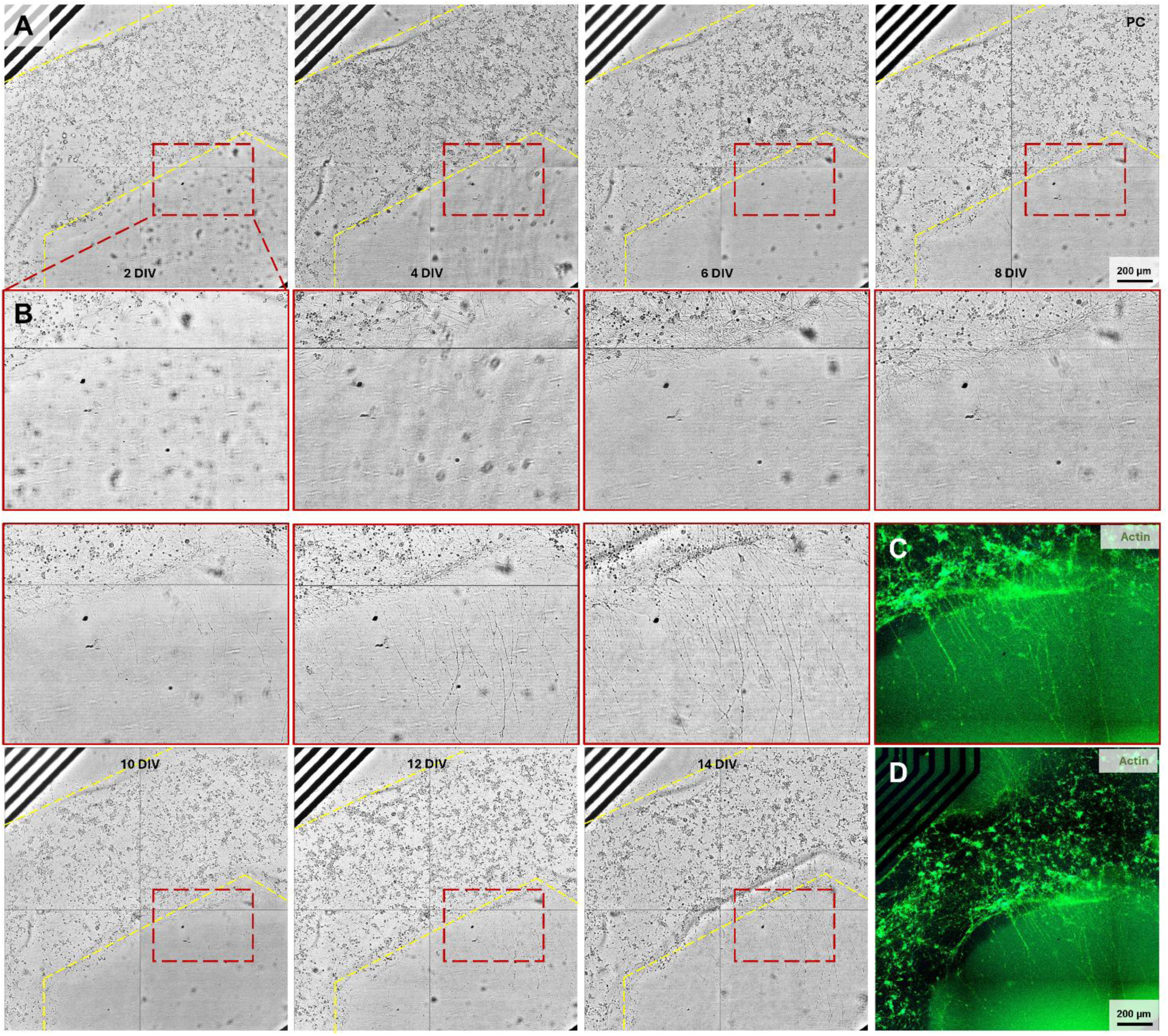
Hydrogel topographies promote self-organisation and neurite extension within engineered microenvironments: (A) representative brightfield time-lapse images showing progressive spatial organisation of the neuronal–glial triculture within the hexagonal GelMA hydrogel architecture. Cells gradually redistributed according to the predefined hydrogel channels, where localised cellular clusters formed during long-term culture; (B) Higher-magnification images reveal elongated neurite-like processes extending from the cell-rich regions into the hydrogel microenvironment; (C) and (D) live-cell actin staining (green) demonstrating large-area neuronal and cytoskeletal organisation throughout the hexagonal hydrogel architecture.

Compared with the hemispherical configuration, the hexagonal grid showed more evidence of forming neurite-like process extensions into the hydrogel region. Elongated cellular processes projecting from cell-dense regions towards and into the patterned hydrogel microenvironment. These results indicate that the hexagonal hydrogel architecture not only influenced cellular distribution but also provide a microenvironment for neurite-like outgrowth inside the gel (Figs. 7B–D). Live-cell actin green staining further demonstrated the spatial organisation of cellular processes within the hexagonal hydrogel microenvironment. Actin-positive structures were distributed throughout the patterned culture and extended into adjacent hydrogel. Actin projections extended beyond the primary cell-dense regions and penetrated the hydrogel. Combined with the brightfield observations, the results suggest that the engineered hydrogel architecture supports cellular process extension beyond the initial regions of cellular aggregation.

Live-cell tubulin red staining confirmed the development of extensive neuronal structures associated with the hexagonal grid architecture. A highly interconnected neuronal-glial network was observed in the whole well and distributed throughout the patterned culture area (Fig. 8). Tubulin-positive neurite-like projections extended from cellular cluster and continuously formed a network spanning neighbouring regions of grids. Neuronal processes penetrate and extend across many millimetres of the hydrogel microenvironment. These observations indicate that neuronal-glial structures were not restricted to the hexagonal channels but were able to extend into the surrounding engineered microenvironment owing to the suitable micromechanics (Figs. 8C, D, G and H). This indicates the potential of engineered photopatterned hydrogels in creating spatially organised neuronal-glial systems with controllable structural complexity. It is also a critical step in investigating how geometry influences network assembly.

**Figure 8.**
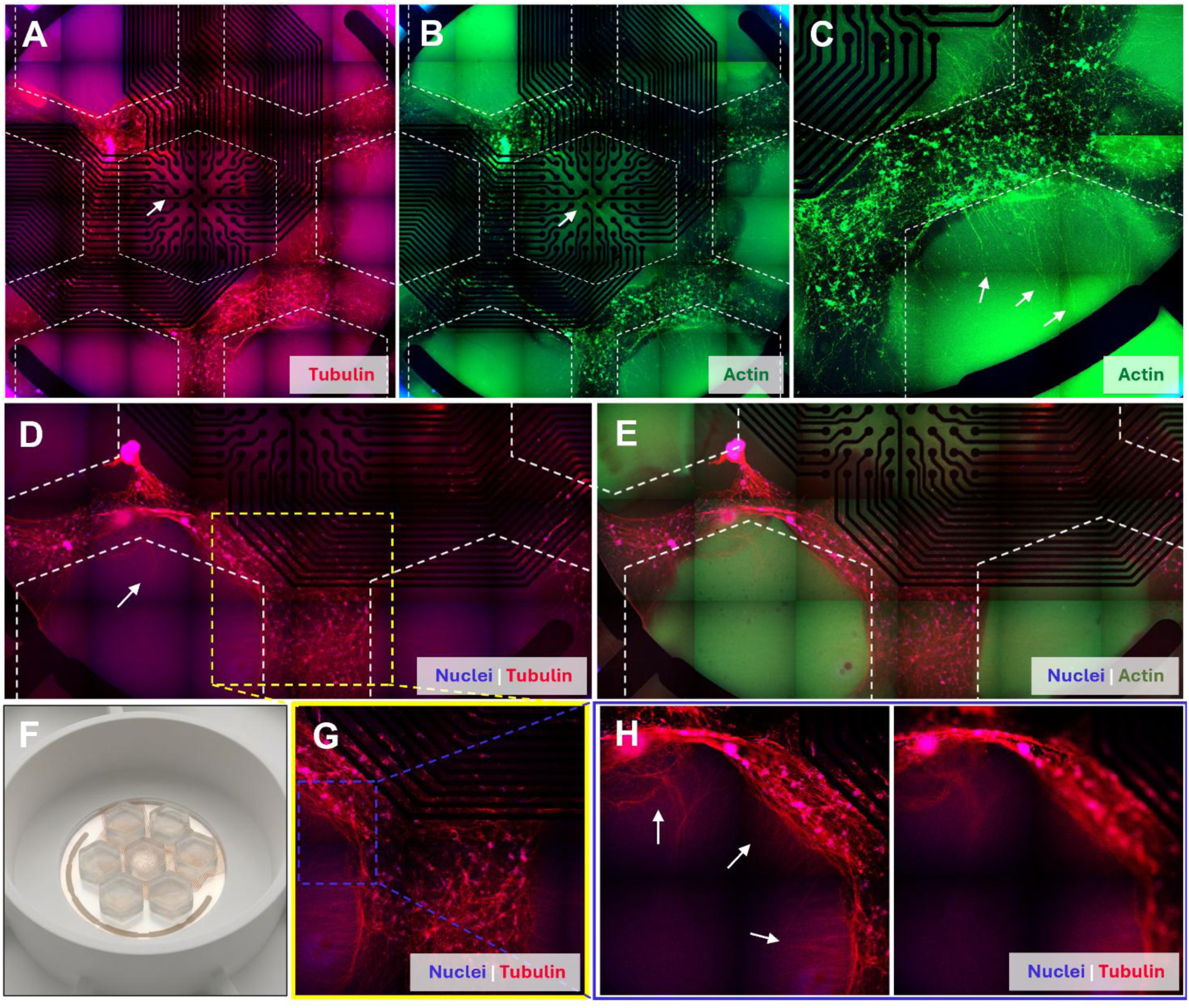
Hydrogel topographies promote self-organisation and neurite extension within engineered microenvironments: (A) whole-well live-cell tubulin staining (red) and (B) actin staining (green) demonstrating large-area neuronal and cytoskeletal organisation throughout the hexagonal hydrogel architecture (C); (D) and (E) representative higher-magnification images showing tubulin-positive neuronal networks and actin-positive cytoskeletal organisation closely associated with engineered hydrogel channels (geometry shown in F); (G) and (H) high-magnification images demonstrating neurite and actin-rich cellular process extension across the hydrogel boundary and into the hydrogel microenvironment.

### 2.6. Neuronal-glial morphology in 3D hydrogel microenvironments

After establishing spatially organised neuronal-glial cultures across engineered hydrogel topographies and observing neuronal extension into the 3D hydrogel microenvironment, we further investigated whether the triculture systems could self-organise within 3D ECMs. Therefore, glutamatergic neurons, GABAergic neurons and astrocytes were embedded within Geltrex and GelMA and cultured under the identical differentiation conditions. Cellular organisation was observed to evaluate network formation within the 3D microenvironment.

Live imaging showed the formation of numerous multicellular aggregates distributed throughout the 3D matrix (Fig. 9A). Unlike the continuous dense network observed in 2D cultures, cells self-organised into discrete clusters interconnected by elongated cellular processes. A presence of a high density of projections extending between neighbouring aggregates, forming a network-like structure in the gel system. These results suggest that cells spontaneously self-organise within 3D matrix and can establish long-range structural connectivity between cellular assemblies (Fig. 9B) and between these assemblies and the matrix boundaries (Fig. 9C).

**Figure 9.**
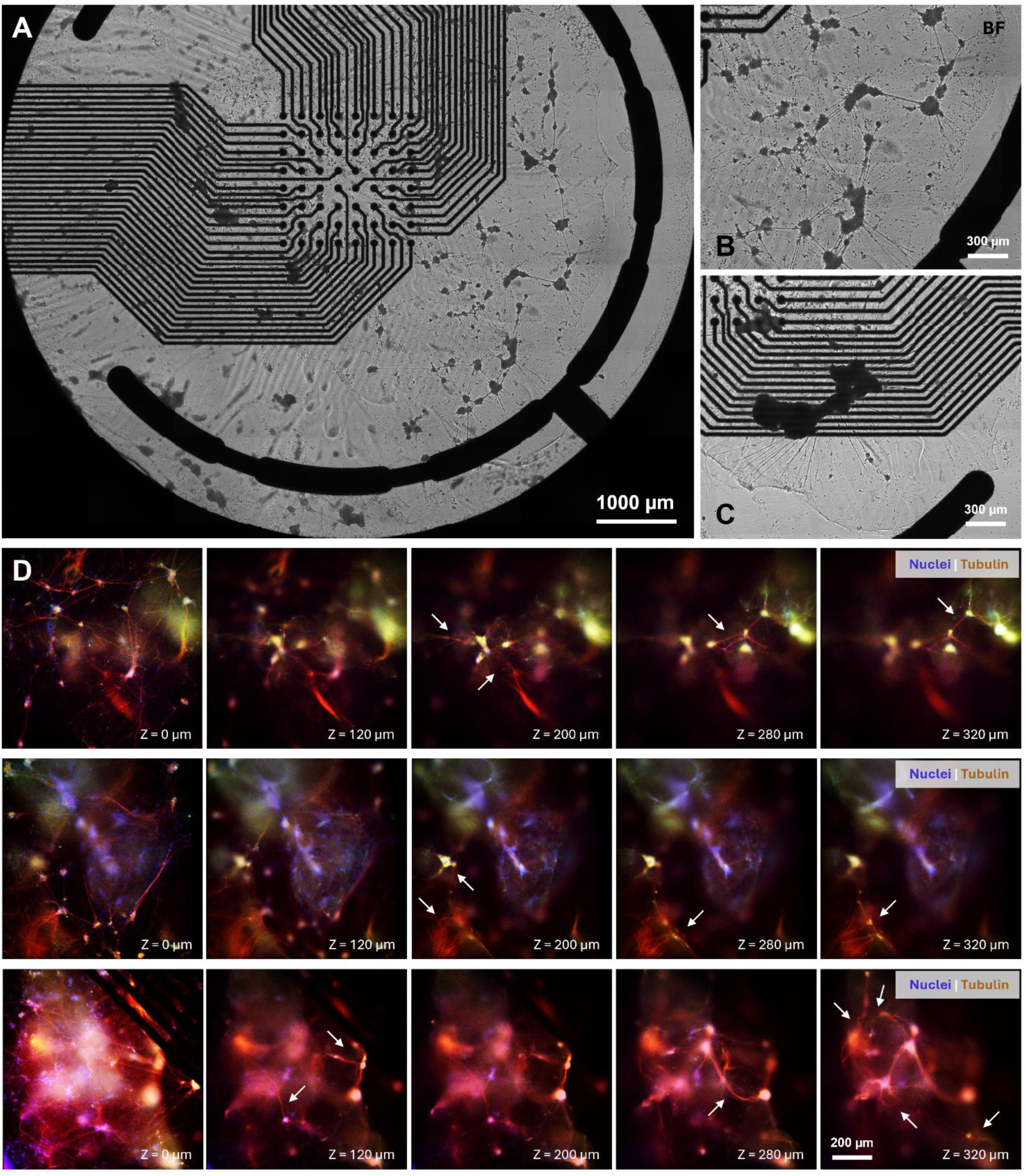
Neuronal–glial self-organisation within a three-dimensional GelMA and Geltrex microenvironment; (A) brightfield image showing the overall morphology of the triculture embedded within the 3D GelMA matrix; (B) and (C) representative higher-magnification brightfield images demonstrating the formation of multicellular clusters interconnected by elongated cellular projections throughout the hydrogel; (D) whole-gel live-cell tubulin staining (red) showing the development of an interconnected neuronal–glial network within the 3D matrix. Confocal z-stack images of three representative regions showing nuclei (blue) and live-cell tubulin staining (red) across the height of the sample in this case Geltrex. Tubulin-positive neurites and neuronal clusters are detected throughout the volume, demonstrating 3D network organisation within the Geltrex matrix.

The neuroglial clustering into organoid-like hubs present in the various configurations reported here, is a survival strategy. Large clusters provide a protective microenvironment where local concentrations of secreted neurotrophic factors such as brain-derived neurotrophic factor (BDNF) and glial cell-derived neurotrophic factor (GDNF), are maintained at higher levels than in sparse cultures^35^. In human iPSC-derived cultures, these hubs represent dense processing centres where glutamatergic and GABAergic neurons form the first functional synapses, often modulated by the accompanying astrocytes^36^.

Neurons are known to be highly mechanosensitive, though their preferences differ by type. PNS neurons such as those from the dorsal root ganglia, typically show maximal outgrowth on substrates with a Young’s modulus of approximately 1 kPa^37^. In contrast, CNS neurons including human hippocampal or cortical iPSC-derived neurons, often show neurite outgrowth that is less dependent on stiffness but highly sensitive to the stability of matrix tethering^37^. The observed preference for growing at the GelMA and Geltrex interface suggests that this provides a high-traction surface that growth cones can exploit to generate the significant mechanical tension required for long-distance elongation^38^. Growth cones sense substrate stiffness by exerting traction forces; these forces correlate with the density of focal adhesions and the strength of the cytoskeletal-substrate coupling^38^. By occupying the interface, the neurite gains the mechanical leverage while remaining bathed in the biochemical cues (laminins and collagens from Geltrex) and trophic support of the overlying GelMA hydrogel^39^.

The fluorescence imaging further confirmed the development of neuronal-glial structures within the 3D microenvironment. Tubulin red positive cellular projections extended from neuronal clusters and formed interconnected networks throughout the matrix (Fig. 9D). The neurite-like structures were observed spanning between neighbouring cellular aggregates, indicating the establishment of structural communication pathways within the 3D microenvironment. Nuclear blue and tubulin-positive red were observed at multiple focal depths across the imaging volume. Cellular aggregates were visible throughout the imaging planes extending to approximately 320 µm, indicating that the neuronal-glial assemblies distributed in the 3D matrix rather than being confined to a single focal plane. Importantly, tubulin-positive projections can be found across multiple depths, suggesting neuronal processes extend within the gel forming 3D networks across the microenvironment.

The transition from 2D topographical cultures to a fully 3D microenvironment resulted in a distinct neuronal-glial organisation. In 2D systems, networks were spatially constrained associated with engineered hydrogel architectures. However, cells embedded within the 3D matrix self–organise into multicellular clusters that connected through extensive neurite-like projections. Similar cluster-based organisation has been reported in neural spheroid and organoid systems, yet our model enables *in situ* differentiation, clustering and formation of interconnected neuronal-glial assemblies directly in 3D biomimetic ECMs from single cell state. These results demonstrate that the neuronal-glial triculture system can self-organise within 3D microenvironments, providing a model for investigating the development of human neuronal-glial micro-circuitry *in vitro*.

For a neurite to extend through or under a crosslinked GelMA network, it must actively remodel the extracellular matrix. Neurites and their growth cones secrete matrix metalloproteinases (MMPs) and serine proteases that degrade target domains of the hydrogel^40^. In fibrin and GelMA gels, neurite extension is significantly enhanced. The patterned domains showcase the morphological significance of this effective degradation and neurite extension. Cells seeded onto the patterns do not simply form isolated organoids; they extend dense, parallel bundles of neurites that trace the geometries connecting multiple organoids across a large area. These neurite bundles exhibit long-term structural stability and are capable of extending continuous, uninterrupted projections over 1-2 mm across the gel, successfully bridging organoids scattered in photopatterned segmented zones. This long-range connectivity is absent in traditional, 2D or 3D cultures, where axons branch randomly and form short, localised connections, failing to establish organised inter-regional pathways^41^.

## 3. Materials and Methods

3.1. **Preparation of media and reagents**

Basal co-culture medium (b:GN) was prepared using Neurobasal medium supplemented with 1× GlutaMAX, 1× B27, and 2-mercaptoethanol. For the induction of precision-reprogrammed neurons, complete glutamatergic neuron medium (comp:GN) was formulated by supplementing b:GN with 10 ng/mL human NT3, 5 ng/mL human BDNF, and 1 µg/mL doxycycline. To facilitate neuronal maturation, 10 µM DAPT was added to the complete medium (comp:GN+D+DAPT) for the 48-hour treatment window beginning on day two. Long-term maintenance and electrophysiological recordings were conducted in BrainPhys complete medium, consisting of BrainPhys basal medium supplemented with 1× B27, 10 ng/mL NT3, and 5 ng/mL BDNF.

### 3.2. Multielectrode array plate coating process

Multielectrode array (MEA) plates (6-well format CytoView, Axion BioSystems) were prepared using a multi-step coating procedure. The MEA plates were sequentially coated with poly-D-lysine (PDL) and Geltrex to promote cell adhesion and neuronal maturation. A vial of PDL (Sigma-Aldrich, D9891) was dissolved in 50 mL of sterile 1x borate buffer to obtain a final concentration of 100 µg/mL. Each well was coated with 350 µL of PDL solution and incubated overnight at 37°C with 5% CO_2_. The following day, wells were rinsed three times with sterile distilled water and air dried for approximately one hour. Geltrex (Thermo-Fisher Scientific) was transferred from −20°C storage to 4°C storage overnight. On the following day, it was thawed on ice for 30 minutes and diluted 1:100 in ice-cold DMEM/F-12. Each well was coated with 350 µL of the diluted Geltrex solution and incubated for 1 hour at 37°C with 5% CO_2_. Excess Geltrex was gently aspirated prior to cell seeding. The 3D photoactive extracellular matrix precursors were always introduced and photopatterned prior to Geltrex coating.

### 3.3. Immunocytochemistry staining of neuronal-glial tricultures on MEA plates

Immunocytochemistry was performed by carefully fixing the neuronal-glial tricultures to minimise disruption of the cell layer and constructs. The culture medium was gently aspirated, and cells were washed once with 1 mL Dulbecco’s phosphate-buffered saline (DPBS) per well for 5 minutes at room temperature. The wash solution was then removed, and 1 mL cold (2–8 °C) 4% paraformaldehyde (PFA) in DPBS was added to each well. Cells were fixed for 10 minutes at room temperature. Following fixation, wells were washed twice with DPBS. For each wash, 1 mL DPBS was added and incubated for 5 minutes at room temperature, after which 900 µL was carefully removed without disturbing the cell layer, leaving a small residual volume in the well. Cells were either processed immediately for staining or stored in 1 mL DPBS at 4 °C with the plate sealed using parafilm for up to two weeks prior to immunostaining. Cells were blocked and permeabilised using a solution consisting of 0.1% Triton X-100 in DPBS supplemented with 3% donkey serum and 3% goat serum. Goat serum and donkey serum were sterile-filtered through a 0.22 µm filter prior to use. Blocking solution was prepared by adding 28.2 mL of 0.1% Triton X-100/DPBS, 900 µL donkey serum, and 900 µL goat serum to a 50 mL centrifuge tube and mixing gently. DPBS was carefully removed from each well, and 1 mL blocking solution was added down the side of the well. Constructs were then incubated for 1 hour at room temperature.

Primary antibodies were prepared in base antibody solution consisting of DPBS supplemented with 10% blocking solution. Per 6-well plate, 6 mL primary antibody solution was prepared by combining 5.4 mL DPBS with 600 µL blocking solution, followed by addition of 12 µL anti-VGLUT2 antibody (final dilution 1:500) and 3 µL anti-MAP2 antibody (final dilution 1:2000). Following the blocking step, the solution was aspirated and 1 mL primary antibody solution was added per well. Plates were sealed with parafilm and incubated overnight at 4°C. After primary antibody incubation, wells were washed three times with DPBS. For each wash, 1 mL DPBS was added, and cells were incubated for 5 minutes at room temperature. Wash solution was carefully removed after each step to avoid disturbing the neuronal layer.

Secondary antibodies were prepared in base antibody solution consisting of DPBS supplemented with 10% blocking solution. Per 6-well plate, 6 mL base antibody solution was prepared from 5.4 mL DPBS and 600 µL blocking solution. 6 µL of Donkey anti-mouse Alexa Fluor 488 (final dilution 1:1000), 6 µL of goat anti-chicken Alexa Fluor 647 (final dilution 1:1000), and 12 µL of DAPI (final dilution 1:500) were then added. DPBS was removed and 1 mL secondary antibody solution was added to each well. Constructs were then incubated for 1 hour at room temperature protected from light. Following secondary antibody incubation, the constructs were washed three times with DPBS. For each wash, 1 mL DPBS was added and incubated for 5 minutes at room temperature, and plates were protected from light throughout. After the final wash, cells were left in DPBS and imaged using a fluorescence microscope with appropriate filters for Alexa Fluor 488, Alexa Fluor 647, and DAPI.

### 3.4. Live-cell staining of neuronal-glial tricultures on MEA plates

Live-cell staining was performed to visualise cytoskeletal organisation and nuclear distribution in neuronal tricultures cultured on MEA plates. Tubulin Tracker™ Deep Red, CellMask™ Actin Green, and Nuclear Blue live-cell stains (Thermo-Fisher Scientific) were prepared in Cell Imaging solution. Prior to staining, culture medium was carefully removed, and cells were washed twice with Cell Imaging solution. Staining solution was prepared in Cell Imaging solution containing Tubulin Tracker Deep Red (1:2000), CellMask Actin Green (1:2000), and Nuclear Blue (1:4000). These dilutions were empirically optimised to minimise cytotoxicity while maintaining sufficient fluorescence signal in neuronal tricultures. Cells were incubated with staining solution at 37°C in a humidified incubator with 5% CO₂ for 40 minutes protected from light. Following incubation, cells were washed twice with Cell Imaging Solution to remove excess dye. Cells were then maintained in 500 µL Cell Imaging Solution per well and imaged immediately using appropriate fluorescence channels.

### 3.5. Establishing a human iPSC-derived glutamatergic and GABAergic neuron, and astrocyte 2D triculture

Cryopreserved human iPSC-derived glutamatergic neurons, GABAergic neurons, and astrocytes (BitBio) were thawed in b:GN medium. Cells were resuspended in pre-warmed complete GN+D medium and counted using trypan blue exclusion with an automated cell counter. Desired cell densities and triculture compositions were prepared per 2D or 3D experiment. The glutamatergic neurons were seeded at a density of 2.8×10^6^ cells/mL, the GABAergic neurons were seeded at a density of 0.9×10^6^ cells/mL, and the astrocytes were seeded at a density ranging from 0.6 ×10^6^ to 1.2×10^6^ cells/mL. The mixed suspensions were adjusted to a final seeding volume of 250 µL per well and carefully seeded into the system. Plates were incubated at 37°C with 5% CO₂. At 24 hours post-seeding, 1.25 mL of fresh, pre-warmed GN+D medium was added gently to each well.

### 3.6. Establishing a human iPSC-derived glutamatergic and GABAergic neuron, and astrocyte 3D triculture

To establish a 3D system, the same ratio as 2D was maintained, and triculture compositions were resuspended in pre-warmed complete GN+D medium (GN+D) and mixed with the desired volume of photoactive extracellular matrix precursor to generate the final mixture. This final composition was gently mixed to ensure homogeneous cell distribution. A desired volume of cell–matrix suspension was seeded per well. Immediately following seeding, 1.25 mL of pre-warmed GN+D medium was added to each well to support cell survival and matrix hydration.

### 3.7. Maintenance of 2D and 3D cultures

Following seeding, cultures in both 2D and 3D conditions were maintained under 37°C and 5% CO₂ humidified incubation environments. At 24 hours post-seeding, 1.25 mL of fresh pre-warmed COMP:GN+D medium was gently added to topographical or 3D culture wells to minimise disturbance of the developing neuronal network. At 48 hours post-seeding, a full medium change was performed with 1.5 mL of GN+D medium supplemented with 10 µM DAPT. On day four, another complete medium exchange was conducted using 1.5 mL/well of GN medium. From day six onwards, cultures were maintained by performing 50% medium changes with BrainPhys complete medium every 48–72 hours (removing and replacing 750 to 1000 µL per well). All media changes were carried out carefully to minimise mechanical disturbance and preserve neuronal connectivity.

### 3.8. Biomaterials and photoactive 3D extracellular matrices

For the photoactive hydrogel matrices, a range of compositions were evaluated. All precursors were prepared in Neurobasal^TM^ growth medium to minimise post-print swelling and transport. Methacrylated gelatin (GelMA) was used at concentrations ranging from 4 to 8% w/v in Neurobasal growth medium. For characterisation purposes, each formulation was photocrosslinked in 24-well cell culture inserts (Merck Millipore) in presence of 0.1% w/v Lithium phenyl-2,4,6-trimethylbenzoylphosphinate (LAP) as the photoinitiator and under configured photoexposure conditions. GelMA was synthesised by dissolving type A porcine gelatin (300 bloom strength) in a 0.25 M carbonate-bicarbonate (CB) buffer at a concentration of 10% (w/v) by stirring at 50°C. The pH of the CB buffer was pre-adjusted to and maintained at 9.0 to optimize the functionalisation reaction. Once the gelatin was completely dissolved, methacrylic anhydride (MAA) was added dropwise to the solution at a ratio of 0.1 mL per gram of gelatin, and the reaction was allowed to proceed for two hours at 50°C with continuous stirring. The reaction was terminated by adjusting the pH of the solution to 7.4. To remove unreacted MAA and other small-molecule impurities, the resulting GelMA solution was dialysed against deionized water at 40-50°C for three days. The purified GelMA solution was then flash-frozen in liquid nitrogen and lyophilised for 72 hours to yield a dry, porous foam, which was stored at −20°C until further use.

Hydrogels were incubated at 37°C in Neurobasal™ medium, minus phenol red (Thermo-Fisher Scientific). To monitor changes in pH during hydrogel incubation, pH measurements were performed at defined time points. At each time point (day 3, 7 and 10), the culture medium from three replicate wells containing the same hydrogel formulation was pooled to provide sufficient volume for accurate pH measurement. Measurements were performed immediately after pooling to minimise CO_2_ exchange. At each designated point (days 1, 3, 7, and 10), hydrogel discs were gently removed from the 24-well Millipore cell culture inserts. The diameter and height of the hydrogels were quantified using ImageJ software (NIH, Bethesda, MD). For each formulation and time point, three replicate hydrogels were analysed, and the mean and SD values of these measurements are reported.

### 3.9. Photo-rheo-Raman measurements

The viscoelastic properties of the hydrogels were evaluated using an Anton Paar photorheometer equipped with a parallel plate geometry and a white light source filtered with a 400-410 nm cutoff. The rheometer gap size was set to 500 μm for all measurements. Oscillatory shear tests were performed at a strain amplitude (γ) of 10% and an angular frequency (ω) of 1 rad/s at a Peltier controlled temperature of 40 °C. The system was equipped with an Anton Paar Cora 5001 Raman spectrometer with a probe positioned towards the material from below the photorheometer glass window to collect Raman data in tandem with photo-exposure and rheology. For each hydrogel formulation, three independent samples were tested, and the results are reported as the mean of the three replicates. *In situ* Raman spectroscopy was employed to monitor the photopolymerisation kinetics of the acrylate system by tracking the intensity variations within the 1600 to 1650 cm^-1^ region. The evolution of the spectra over the exposure timeline was monitored to verify the relative stability of the primary spectral features. To quantify the photopolymerisation progress, the fractional conversion α_t_ was determined by tracking the intensity of the aliphatic C=C double bond stretch of the monomer at 1644 cm^-1^ over time. The spectra were baseline corrected and the intensity of this reactive band I_1644_ was normalised relative to an invariant internal reference peak located at 1614 cm^-1^ (I_1614_), and this ratio was used for conversion calculations.

## 4. Conclusions

The developed 3D human neuroglial triculture platform represents a substantial advancement over conventional 2D *in vitro* models and classical unguided 3D organoid systems. Combining deterministic cellular reprogramming with a mechanically compliant, photopatterned hydrogel system, the platform successfully recapitulates both the structural complexity and functional dynamics of native neural circuits. The experimental findings support several translationally relevant conclusions. Utilising low-concentration GelMA gels, suppresses reactive gliosis and permits robust, 3D volumetric network arborisation. Replacing heterogeneous, slow-maturing directed differentiation protocols with deterministic cell populations ensures standardised and highly reproducible cellular starting points, which are essential for high-throughput pharma screening and disease modelling.

Photopatterned surface topographies provide critical contact guidance cues that orient and align clusters and guide long-range neurite projections. Excitatory monocultures spontaneously default to pathological, highly synchronised burst-suppression patterns. The inclusion of standardised populations of inhibitory GABAergic interneurons and supporting astrocytes is required to establish physiological, temporally diverse, and complex network-level firing dynamics. Advanced computational metrics computed from high-density MEA recordings including entropy, complexity, and manifold volume, provide highly sensitive, non-invasive biomarkers of network maturation and functional complexity that go beyond conventional, linear firing rate analyses. In conclusion, this integrated bioengineering approach establishes a highly robust, controllable, and translationally relevant *in vitro* platform for investigating neurodevelopmental programs, mapping inter-regional axonal pathfinding, and screening therapeutic compounds targeting complex human neural networks.

Apart from the functional and electrophysiological characteristics of the model, this study further demonstrates the importance of engineered spatial configuration in regulating neuronal-glial organisation. Photopatterned hydrogel topographies spatially navigate cellular assemblies, in this case controlling the network geometry at the millimetre scale. Hemispherical grids promote boundary-associated cluster formation, and hexagonal grids with vertical walls support extensive neurite guidance as conduits and extension into 3D hydrogel environment. These results indicate that engineered hydrogel architectures can be used not only as cell-supporting matrices, but also as navigators of cellular network organisation. Additionally, triculture systems were embedded in 3D matrices showing the spontaneous self-organisation of interconnected neuronal-glial assemblies throughout the ECM. Unlike conventional organoid systems that depend on fusion of pre-differentiated cellular aggregates, this platform supports *in situ* differentiation, cellular organisation and network formation within the 3D matrices. This finding indicates a promising model for studying human neural circuit formation and dysfunction *in vitro*, giving the opportunity for the development of neurodegeneration diseases and drug discovery.

The integration of multi cell type cellular organisation, matrix engineering, photopatterned biofabrication and electrophysiological interrogation establishes a versatile platform for constructing human neuronal-glial systems. This platform provides substantial potential for modelling neurodevelopmental and neurodegenerative diseases, investigating the mechanisms of network assembly and decline, and exploring novel therapeutic interventions. Future work will focus on extending the platform to vascularised and neurovascular models, which can achieve further organoid network-level insights on human brain physiology and pathology *in vitro*.

## 5. Author contributions

H.H. conceived the project and designed the research. S.D. and H.H. developed the experimental plan for tissue modelling and biofabrication and performed the experimental cell culture work. H.H., S.D., and D.W. formulated the biomaterials and performed biomaterials screening. H.H. performed the image and biological data analysis. All authors were involved in writing the manuscript text.

## Acknowledgements

The authors would like to thank the UK Research and Innovation (UKRI). This work was supported by the UKRI Future Leaders Fellowship [grant number MR/X034976/1]

## 7. Data availability

The images experimentally acquired and analysed during the current study are available from the corresponding author on reasonable request.

