## Supplementary Information for "Photopatterned spatiotemporal organisation and *in situ* differentiation of 3D human cortical networks"

**Submitted for consideration by Advanced Science**

**Authors:** Siyuan Dong[1], David Weyland[1], Hossein Heidari[1]*

[1] *Institute for Materials Discovery, University College London, London E20 2AE, UK*

### 1. Photopolymerisation kinetics and mechanics

Across all formulations (6% to 10% w/v), the storage modulus (G') remains significantly higher than the loss modulus (G''), confirming the formation of robust, covalently crosslinked hydrogel networks with predominantly elastic behaviour. The storage modulus exhibits an exponential increase, rising from approximately 65 Pa at 6% w/v to over 1,200 Pa as the concentration reaches 9–10% w/v. This substantial increase in G' is attributed to the higher crosslinking density and reduced mesh size within the hydrogel matrix at higher polymer fractions. The loss modulus (G'') shows a minor, steady increase from 0.3 Pa to roughly 3 Pa, reflecting slight increases in viscous dissipation within denser networks. Crucially, the crossover modulus remains consistently low (~0.2 Pa) across all concentrations, indicating that the initial transition state stiffness is relatively independent of the ultimate network density.

The real-time *in situ* photorheological profile of 6% w/v GelMA hydrogel tracks structural evolution as a function of applied energy dosage ($mJ/cm^3$). At low energy dosages (< 30 $mJ/cm^3$), the sample behaves as a viscoelastic liquid where the loss modulus (G'') sits slightly above or equal to the storage modulus (G'). A distinct crossover point is observed at an energy dosage of approximately 30 $mJ/cm^3$, marking the formal gelation threshold where a continuous, space-spanning macromolecular network is established. G' rises sharply by several orders of magnitude, capturing the rapid propagation of covalent crosslinks. The slope of G' begins to mature and plateau near 40–50 Pa as the energy dosage approaches 120 $mJ/cm^3$, implying that the available methacryloyl groups are nearing maximum radical conversion under these curing conditions.

Dissolved oxygen acts as a radical scavenger, forming unreactive radicals that halt chain propagation. Higher concentrations of GelMA introduce a higher density of methacryloyl groups per unit volume. This increased local concentration of reactive double bonds increases the probability of radical-monomer collisions, allowing the system to consume dissolved oxygen more rapidly and initiate network formation sooner. The *in situ* monitoring of the acrylate photopolymerisation kinetics yielded an initial rapid alteration in the normalised peak ratio within the first 90 s of exposure, reaching a calculated conversion variation of 23.63%. Beyond this initial regime, the system transitioned into a steady-state characteristic of a non-reactive spectator environment. Non-linear regression using a first-order kinetic model applied to the data yielded an asymptotic plateau amplitude (A) of -26.59% and an apparent reaction rate constant (k) of 0.00297 $s^{-1}$ (corresponding to an apparent kinetic time constant $\tau$ of 337 s).

### 2. Excitatory-inhibitory balance and maturation

A critical limitation of single organoid and also monoculture or biculture neural networks is a lack of physiological feedback loops. When hiPSC-derived excitatory glutamatergic neurons are cultured in isolation on standard 2D substrates, they spontaneously generate highly synchronised, repetitive, and stereotypic firing patterns known as identical burst suppression (IBS). This behaviour is characterised by abrupt, explosive bursts of electrical activity followed by periods of complete silent suppression, a clinical signature that directly mirrors the pathological brain states observed in severe postanoxic coma and deep anaesthesia. The developed platform resolves this pathological phenotype by establishing a precise excitatory-inhibitory balance. By coculturing glutamatergic neurons with GABAergic inhibitory interneurons and hiPSC-derived astrocytes, the platform introduces physiological feedback loops.

The mature GABAergic neurons release the inhibitory neurotransmitter GABA, which acts on post-synaptic $GABA_A$ receptors to hyperpolarise the membrane and suppress runaway excitation. This active inhibition modulates the network, converting the pathological, highly synchronised IBS bursts into complex, asynchronous, and temporally diverse firing patterns. Astrocytes actively participate in excitatory-inhibitory regulation by expressing glutamate transporters (such as GLT-1/EAAT2), which rapidly clear excess glutamate from the synaptic cleft, preventing excitotoxicity and maintaining stable, long-term network excitability. As the triculture matures within the 6% GelMA hydrogel, the network undergoes a highly predictable functional evolution. Spontaneous firing is initiated early, and as synapses form and refine, the network achieves stable, coordinated activity by DIV 25. This mature state is sustained over long-term culture, showing progressive improvements in key parameters. Weighted mean firing rate steadily increases and stabilises, reflecting robust individual neuronal excitability and active synaptic transmission. Burst frequency and duration transitions from long, unstructured, and infrequent events to highly controlled, short, and frequent network bursts, indicating the refinement of synaptic connections.

#### 3. Cell adhesion and spreading on hydrogel substrates and surfaces

Live imaging of the platform captured sequentially over multiple weeks, provides a detailed view of the dynamic self-assembly process. At 0 DIV, cells are observed as a homogeneous, isolated population of single-cells scattered across the photopatterned topographies. By 1 DIV and 2 DIV, cells begin to actively navigate the micropatterns, exhibiting local migration and alignment along the pre-defined geometries. Over the subsequent days (7-12 DIV), the cells do not remain static; they exploit the cell-adhesive RGD peptide sequences and compliant mechanics of the 6% GelMA matrix, actively migrating toward the intersection nodes of the patterned geometries. By 14 DIV and 16 DIV, this localised migration culminates in the self-assembly of cells into highly dense, spherical multicellular aggregates. Comparisons between 0 DIV and 18 DIV demonstrate that the cells have clustered into stable, spherical mini-organoids or assembloids containing thousands of cells. Importantly, these self-assembled aggregates are connected to adjacent clusters by thick, highly stable, and parallel neurite cables that trace the permissive patterned pathways, establishing a highly organised, closed-loop circuit across the hydrogel surface.

#### 4. Medium compositions and reagents

The supplementary table (S1) shows the reagent composition for various media and cocktails used for cell culture development and staining.

**Table S1. Information on reagents, cell lines and kits used in this study**

| **REAGENT or RESOURCE** | **SOURCE** | **IDENTIFIER** |
|---|---|---|
| Chemicals, medium, antibody and recombinant proteins | | |
| Geltrex™ LDEV-Free Reduced Growth Factor Basement Membrane Matrix | Fisher Scientific | Cat# A1413202 |
| Poly-ᴅ-Lysine Hydrobromide | Sigma | Cat# P6407 |
| Borate buffer (20X) | Fisher Scientific | Cat# 28341 |
| Sterile water | Sigma | Cat# W3500 |
| DMEM/F-12, HEPES | Fisher Scientific | Cat# 11330032 |
| Neurobasal™ Medium | Fisher Scientific | Cat# 21103049 |
| B-27™ Supplement (50X), serum free | Fisher Scientific | Cat# 17504044 |
| GlutaMAX™ Supplement | Fisher Scientific | Cat# 35050061 |
| 2-Mercaptoethanol (50 mM) | Fisher Scientific | Cat# 31350010 |
| Recombinant Human NT-3 Protein | Bio-techne / R&D Systems | Cat# 267-N3-005 |
| Recombinant Human BDNF Protein | Bio-techne / R&D Systems | Cat# 248-BDB-005 |
| DAPT | Bio-techne / R&D Systems | Cat# 2634 |
| Doxycycline hyclate | Sigma | Cat# D9891 |
| Bovine Serum Albumin | Sigma | Cat# A7906 |
| BrainPhys™ Neuronal Medium | Stem Cell Technologies | Cat# 5790 |
| DPBS, no calcium, no magnesium | Fisher Scientific | Cat# 14190144 |
| Pierce™ 16% Formaldehyde (w/v), Methanol-free | Fisher Scientific | Cat# 11586711 |
| Triton™ X-100 | Sigma-Aldrich | Cat# T8787-50ML |
| Donkey serum | Sigma-Aldrich | Cat# D9663 |
| Goat serum | Sigma-Aldrich | Cat# G9023 |
| DAPI | Bio-techne / R&D Systems | Cat# 5748/10 |
| Anti-MAP2 antibody | Abcam | Cat# ab5392 |
| Goat anti-Chicken IgY (H+L) Secondary Antibody, Alexa Fluor™ 647 | Thermo Fisher Scientific | Cat# A-21449 |
| Critical commercial assays | | |
| HCS NuclearMask™ Stains | Fisher Scientific | Cat# H10325 |
| Tubulin Tracker™ Deep Red | Fisher Scientific | Cat# T34077 |
| CellMask™ Green Actin Tracking Stain | Fisher Scientific | Cat# 57243 |
| Experimental models: Cell lines | | |
| Human iPSC-derived glutamatergic neurons | Bit.bio | Cat# io1001 |
| Human iPSC-derived GABAergic neurons | Bit.bio | Cat# io1003S |
| Human iPSC-derived astrocytes | Bit.bio | Cat# ioEA1093 |
